# Dynamics of biofilm-forming *Bacillus subtilis* in *Caenorhabditis elegans* gut

**DOI:** 10.1101/2025.01.26.634914

**Authors:** Yasuaki Saitoh, Tian (Autumn) Qiu

## Abstract

Biofilm-forming *Bacillus subtilis* strain NCIB3610 has been reported to show biofilm-mediated beneficial effects when fed to *Caenorhabditis elegans*, including enhanced stress resistance, extended lifespan, and protection against neurotoxic agents. However, biofilm-forming *B. subtilis* presents multiple cell types, and thus a thorough characterization of *B. subtilis* cell dynamics in the *C. elegans* gut *in vivo* is necessary to understand mechanisms of the biofilm-mediated beneficial effects. In this study, we utilized an isogenic NCIB3610 strain expressing a biofilm reporter P*tapA-mKate2* to monitor matrix-producing components in *C. elegans* gut. Interestingly, our results revealed that in live *C. elegans* gut, no *B. subtilis* biofilms were formed but only free spores and diffused fluorescence, which were likely from dietary matrix-producing cells crushed by the pharynx. Additional data showed that spores can geminate into motile cells but not matrix-producing cells in living *C. elegans* gut. Biofilm formation was found in gut after *C. elegans* death, indicating that mechanistically, life-dependent functions of the worm inhibit the formation of matrix-producing cells in gut. These findings resolved a key piece of puzzle in understanding the fundamental mechanisms of *C. elegans-B. subtilis* interaction and highlighted the importance of characterizing cell dynamics *in vivo* in host-microbe interaction research.

## Background

Biofilms are diverse microbial communities encased in a self-produced extracellular matrix, and are one of the most adaptive forms of bacteria. The extracellular matrix serves to protect cells from external stressors, while also generating close contact between adjacent cells, promoting cell-to-cell communications. Biofilm growth can be concerned to various medical and industrial problems, such as infections from contaminated medical devices, food spoilage, pipe blockages, and metal corrosion in ships (Shineh et al., 2023). Conversely, biofilms formed by rhizosphere bacteria in plants stimulate plant growth and provide biological control against soil-borne pathogens (Hashem et al., 2019). Biofilms also naturally colonize different surfaces of the body, such as the digestive tract, lungs, vagina, and skin (Buret and Allain, 2023). Their presence in these areas contributes to maintaining homeostasis on those mucosal surfaces. Disruption of beneficial biofilm can cause pathogenic bacteria to invade or cause pathogenicity in dispersed bacteria. Thus, biofilms can be both harmful and beneficial.

*Bacillus subtilis*, a non-pathogenic Gram-positive soil bacterium, has been used as a model organism for studying biofilm formation (Vlamakis et al., 2013; Mielich-Süss and Lopez, 2015; Schoenborn et al., 2021; Arnaouteli et al., 2021). Biofilm-forming strains of *B. subtilis* can form robust biofilms, as indicated by highly structured floating pellicles that grow on the surface of liquid cultures and colonies that grow on agar plates. The undomesticated NCIB 3610 strain, with a sequenced genome, is capable of forming robust, highly structured colonies compared to the domesticated laboratory strain 168 and its derivatives. Within *B. subtilis* biofilms, genetically identical cells express different genes to generate subpopulations of functionally distinct co-existing cell types. This process occurs through stages of development, maturation, and disassembly of the community. Motile cells with flagella differentiate into non-motile, matrix-producing cells in response to external signals, and are organized into chains and surrounded by an extracellular matrix. In mature biofilms, part of the motile cells and matrix-producing cells become sporulating cells and form endospores. The released free spores, which possess high resistance to environmental stress, germinate under favorable conditions to resume the motile state.

*Caenorhabditis elegans* is a well-established model for studying host-microbe interactions. *C. elegans* naturally habitat in decomposing plant matter (Kiontke et al., 2011; Félix and Duveau, 2012), where *B. subtilis* and related bacteria multiply (Siala et al., 1974). The complete genome sequence has already been revealed (C. elegans Sequencing Consortium, 1998), and genetic manipulation such as transgenesis and gene knockout can be performed relatively easily. The transparent body of *C. elegans* facilitates visualization of bacterial colonization in its gut, including pharynx, intestine, and rectum (McGhee, 2007). Bacterial food taken from the mouth by pharyngeal pumping is crushed in the pharyngeal grinder, digested in the intestine, passed to the rectum, and expelled through the anus to the outside of the body as defecation. Importantly, a small proportion of the ingested bacteria may survive the crushing and digesting and colonize in the intestine (Vega and Gore, 2017; Dirksen et al., 2020). In the laboratory, the non-pathogenic *Escherichia coli* strain OP50 is used as a standard food source (Brenner, 1974), and this strain lacks the ability to form biofilms (Arata et al., 2020).

Multiple studies have reported beneficial effects of biofilm-forming *B. subtilis* on *C. elegans* (Donato et al., 2017; Smolentseva et al., 2017; Goya et al., 2020; Cogliati et al., 2020). *C. elegans* raised on the NCIB3610 strain as a diet exhibited increased lifespan, enhanced resistance to stress and pathogens, and protection against neurotoxic agents such as α-synuclein and amyloid β, compared to those raised on the OP50 strain. These beneficial effects diminished when feeding isogenic biofilm-deficient mutants, suggesting that biofilm formation is strongly involved in such effects. Despite previous results, there is a lack of evidence on whether *B. subtilis* can form biofilms *in vivo* in the *C. elegans* gut. Understanding the dynamics of biofilm-forming *B. subtilis* in the worm gut is critical for us to elucidate further mechanisms of the beneficial effects by biofilm-forming *B. subtilis*. Herein, we aimed to investigate the dynamics of *B. subtilis* cells in *C. elegans* gut. Using an isogenic *B. subtilis* NCIB3610 strain expressing a fluorescence reporter for biofilm formation, we investigated the status of biofilm formation and compositions of *B. subtilis* cells *in vitro* and in *C. elegans* gut. Our results show that *B. subtilis* NCIB3610 robustly form biofilm *in vitro* but not in *C. elegans* gut. Instead, diffused fluorescence—likely from grinded matrix-producing cells in diet—and free spores were found in the *C. elegans* intestinal lumen, and these contents were readily replaceable by switching to another food. Furthermore, we discovered that biofilm formation only formed after *C. elegans* death. Our results showed that live *C. elegans* gut did not support biofilm formation under the experimental conditions we have tested, and the previously reported beneficial effects of biofilm-forming *B. subtilis* may be attributed to the cell content released from grinded matrix-producing cells instead of live biofilm formation in the *C. elegans* gut.

## Methods

### *C. elegans* and bacterial strains

*C. elegans* strains wild-type Bristol N2, KU25 *pmk-1(km25)* and CF1038 *daf-16(mu86)*, and *E. coli* strains OP50 and OP50-GFP were purchased from the Caenorhabditis Genetic Center (University of Minnesota, Minneapolis, USA). *B. subtilis strain* YCN095 (*sacA*::P*_tapA_*-*mKate2*, kan^R^. Also named KG091) was generously gifted by Prof. Yunrong Chai (Northeastern University, Boston, USA) (Gozzi et al., 2017). Unless otherwise noted, all *C. elegans* strains were maintained at 20°C on nematode growth medium (NGM) agar plates (3 g/L NaCl, 2.5 g/L peptone, 17 g/L Bacto-agar, 5 mg/L cholesterol, 1 mM CaCl_2_, 1 mM MgSO_4_, 25 mM KPO_4_ [pH 6.0]) seeded with OP50 (Brenner, 1974). OP50-GFP and YCN095 bacteria were streaked on lysogeny broth (LB) agar plates (10 g/L Tryptone, 5 g/L yeast extract, 10 g/L NaCl, 15 g/L Bacto-agar) containing 100 µg/mL ampicillin and 50 µg/mL kanamycin, respectively, stored at 4°C and used within one month. OP50-GFP colonies were inoculated into LB liquid and cultured overnight at 37°C with shaking in an orbital shaker. The overnight culture of 100 µL was dropped onto NGM agar in a 60 mm Petri dish, dried, and incubated at room temperature for 24 to 29 hours.

### Formation of YCN095 colony biofilm and pellicle biofilm

For colony biofilm formation, YCN095 colonies were inoculated into 2 mL of LB liquid containing 50 µg/mL kanamycin and cultured overnight at 37°C with shaking in an orbital shaker. The overnight culture of 80 µL was added to 4 mL of LB liquid and cultured for 3 hours at 37°C with shaking. A 2.5 µL sample of the exponential phase culture (OD600 = 0.5–0.6) was dropped onto LB agar or LBGM agar (LB agar containing 1% glycerol and 100 µM MnSO_4_) and incubated at 30°C for 72 hours. For pellicle biofilm formation, 20 µL of the exponential phase culture was inoculated into 2 mL of LB or LBGM liquid in a 24-well plate and cultured statically at 30°C for 72 hours. Colony and pellicle biofilms formed after 48 hours were photographed using an Apple iPhone SE. Fluorescence imaging of P*tapA-mKate2* expression was performed under a Leica MZ10 F fluorescence stereomicroscope equipped with a Zeiss Axiocam 305 mono camera at an exposure time of 50 ms. Contrast and brightness of fluorescence images were adjusted appropriately on Zeiss ZEN 3.7 software with gamma set to 1.0.

### Preparation of YCN095 mixed-cell plates and spore plates

For preparation of YCN095 mixed-cell plate, 100 µL of the exponential phase culture described above was dropped onto NGM agar in a 60 mm Petri dish, dried, and incubated at 20°C. For preparation of YCN095 spore plate, spore preparation and purification were performed as previously described with some modifications (Donato et al., 2017; Goya et al., 2020). YCN095 colonies were inoculated into 20 mL of Schaeffer’s sporulation medium liquid (8 g/L nutrient broth, 13.4 mM KCl, 487 µM MgSO_4_, 1 mM Ca(NO_3_)_2_, 10 µM MnCl_2_, 1 µM FeSO_4_, pH adjusted to 7.6 with NaOH) and cultured at 37°C for 48 hours with shaking. Heat treatment at 80°C for 20 minutes was performed to kill cells but not spores. After centrifugation (4,200 g, 5 min, 15°C), the pellet was treated with a 25 µg/mL lysozyme solution at 37°C for 30 minutes with shaking at 100 rpm in an orbital shaker. After centrifugation (4,200 g, 5 min, 15°C), the pellet was washed with sterile water. This cycle of lysozyme treatment and washing was repeated three times in total. The purified spores were suspended in 1 mL of sterile water, and a portion of the suspension was used to check the purity of the purification under a microscope, while the remaining suspension was dispensed and stored at –80°C. To culture spores, 100 µL of the purified spore suspension was dropped onto NGM agar or peptone-free NGM agar (PF-NGM; 3 g/L NaCl, 17 g/L Bacto-agar, 5 mg/L cholesterol, 1 mM CaCl_2_, 1 mM MgSO_4_, 25 mM KPO_4_ [pH 6.0]) in a 60 mm Petri dish, dried, and then cultured at 20°C for an appropriate period of time.

### Determination of the cell type composition of YCN095 lawns

The whole bacterial lawn from YCN095 mixed-cell plates incubated for 24, 72, 120 and 168 hours, as well as YCN095 spore plates incubated for 24, 48 and 72 hours, was collected using a cell scraper with 3 mL of M9 buffer. The collected suspensions were subjected to a 10-second sonication cycle in an Emerson Branson 2800 ultrasonic bath, followed by thorough pipetting to suspend. This cycle of sonication and pipetting was repeated five times to generate a single-cell suspension. A 0.5 µL aliquot of the suspension was mounted on a 2% thin agarose pad and imaged under a Zeiss Axio Imager.Z2 upright microscope equipped with a Plan-APO 40x/1.4 Oil lens.

For bright-field imaging, the light source intensity and exposure time were set to 4.2 V and 50 ms, respectively. For green fluorescence imaging, the light source intensity, exposure time, excitation wavelength, and emission wavelength were set to 15%, 50 ms, 450–490 nm, and 500–550 nm, respectively. For the red fluorescence imaging, these parameters were set to 15%, 200 ms, 550–580 nm, and 590–650 nm.

Images were acquired at the focal plane where spores appeared dark in the bright field. Contrast and brightness of images for each channel were adjusted appropriately on Zeiss ZEN 3.7 software with gamma set to 1.0. At least 1,000 cells per sample were analyzed for cell type and the presence of P*tapA-mKate2* using NIH ImageJ software. Criteria for cell identification are detailed in the main text. Green channel images were used to distinguish specific P*tapA-mKate2* red fluorescence from autofluorescence.

### Microscopic visualization of bacteria in the gut of *C. elegans*

#### C. elegans cultured on YCN095 mixed-cell plates

Approximately 80 L1 larvae, prepared via the standard alkaline bleach method from *C. elegans* populations maintained on YCN095 mixed-cell plates for several generations, were transferred to a day-5 YCN095 mixed-cell plate (120–125 hours post-inoculation) and cultured at 20°C. The *C. elegans* obtained after 72 hours was defined as adult day 1 worms. Adult worms were grown on YCN095 and transferred to fresh day-5 YCN095 mixed-cell plates at days 1, 3, 5, 7, and 9 using a platinum wire to maintain food conditions and prevent progeny contamination. Fluorescence microscopy was performed on L4 larvae (48 hours post-L1 transfer) and adult worms at days 2, 7 (before transfer to plates with new food), 12, and 16. To analyze worms that had died of old age, all dead adult worms on the plate containing adult day 16 worms were removed, and the plate was cultured for an additional 2 days at 20°C. Newly deceased adult worms were then subjected to fluorescence microscopic analysis.

#### Food switch experiment with OP50-GFP and YCN095

Approximately 80 L1 larvae were prepared from populations maintained on OP50-GFP or YCN095 mixed-cell plates for several generations. The larvae were transferred to day-1 OP50-GFP plates (24–29 hours post-inoculation) or day-5 YCN095 mixed-cell plates and cultured at 20°C. Adult worms at days 1, 3, and 5 were transferred to fresh OP50-GFP or YCN095 plates using a platinum wire to maintain lawn conditions and prevent progeny contamination. Adult day 7 worms cultured on OP50-GFP or YCN095 mixed-cell plates were transferred to the opposite lawn type (day-5 YCN095 or day-1 OP50-GFP) after their surface bacteria were removed by crawling the worms on NGM agar without food for approximately 30–60 seconds. After food switching, worms were cultured for an additional 60 minutes at 20°C before fluorescence microscopy.

#### C. elegans cultured on YCN095 spore plates

Approximately 80 L1 larvae were prepared from *C. elegans* populations maintained for several generations on OP50-GFP plates. These larvae were transferred to a day-1 OP50-GFP plate and cultured at 20°C. On reaching adulthood (day 1), worms were transferred to a freshly prepared YCN095 spore PF-NGM plate using a platinum wire. Prior to transfer, bacteria attached to their bodies were removed by crawling the worms on NGM agar plates without food. The worms were then cultured at 20°C. To maintain the condition of the bacterial lawn and prevent contamination from progeny, adult worms at days 3 and 5 were transferred to freshly prepared YCN095 spore PF-NGM plates using a platinum wire. Fluorescence microscopic analysis was conducted on adult worms at day 6.

#### Fluorescence microscopic analysis

For all fluorescence analyses, worms were mounted in 2 µL of M9 buffer containing 10 mM sodium azide on a 2% thin agarose pad and imaged under a Zeiss Axio Imager.Z2 upright microscope equipped with a Plan-APO 40x/1.4 Oil lens. Imaging settings for the bright-field, red and green fluorescence channels were consistent with those used for the YCN095 cell-type composition analysis, except that the green channel exposure time in the food switch experiment was reduced to 10 ms. Contrast and brightness of each channel image were adjusted appropriately on Zeiss ZEN 3.7 software with gamma set to 1.0. P*tapA-mKate2*-positive spores in the worm gut were identified based on their characteristic bright-field focal image and oval shape (approximately 1 µm in length) in the red channel. Matrix-producing cells were identified as P*tapA-mKate2*-positive rod-shaped cells exceeding 2 µm in length in the red channel. Green channel images, excluding those from the food-switch experiment, were used to distinguish specific red fluorescence of P*tapA-mKate2* from autofluorescence. At least eight worms were analyzed per condition in at least two independent experiments.

### Measuring bacterial cells in *C. elegans* gut

The procedures for body surface sterilization, lysis of *C. elegans*, and colony-forming unit (CFU) assays were conducted following established protocols with minor modifications (Laaberki and Dworkin, 2008; Donato et al., 2017; Dirksen et al., 2020). Approximately 100 L1 larvae were prepared using the standard alkaline bleach method from *C. elegans* populations maintained over several generations on YCN095 mixed-cell plates or OP50 plates. The larvae were then transferred to a day-5 YCN095 mixed-cell plate or a day-1 OP50 plate, respectively, and cultured at 20°C. To maintain the integrity of bacterial lawns and prevent contamination from worm progeny, adult worms at days 2, 3, and 5 were transferred to fresh day-5 YCN095 mixed-cell plates or day-1 OP50 plates after collection and washing five times with M9 buffer. Adult worms at day 2 (prior to transferring to fresh plates) and day 7 were collected in 1.5 mL tubes using M9 buffer and washed five times with M9 buffer containing 0.025% Triton X-100. To close the worms’ mouths, the collected worms were incubated in M9 buffer containing 10 mM levamisole for 1 minute, followed by removal of the supernatant. The worms were then surface sterilized by incubation in M9 buffer containing 4% (v/v) Clorox for 2 minutes, followed by five washes with M9 buffer. Throughout all washing and collection steps, worms were allowed to sink by gravity for approximately 40 seconds rather than using centrifugation to separate the adults from the bacteria and progeny. The washed worm pellet was resuspended in 1 mL of M9 buffer and transferred onto an NGM plate without food. The exact number of worms was counted under a stereomicroscope. All worms on the NGM plate were then transferred to a Benchmark Scientific prefilled 2 mL tube with 1.5 mm diameter zirconia beads using a Pasteur pipette to avoid worm loss. After allowing the sample to stand for 1 minute to let the worms settle, a portion of the supernatant was sampled as a blank for the CFU assay, and the remaining supernatant was removed to prepare a 500 µL worm suspension. The worms were lysed completely by processing for 170 seconds at speed 1 using a Fisherbrand Bead Mill 4 Mini Homogenizer.

For worms cultured on YCN095 mixed-cell lawns, 200 µL of the worm lysate was heat-treated at 80°C for 20 minutes to kill all cells except for spores. Serial dilutions from heat-treated and untreated worm lysates of 50 µL were plated on LB agar and incubated at 37°C for 24 hours. The resulting colony counts were used to calculate CFU per worm. The CFU per worm derived from heat-treated lysates represented the number of spores, while the difference between CFU from untreated lysates and heat-treated lysates represented the number of motile cells. For worms cultured on OP50 lawns, CFU per worm was determined from untreated lysate dilutions. Colony formation from undiluted blank samples was verified to be either zero or negligible.

### Statistical analysis

All data were analyzed for normality and variance using Shapiro–Wilk test and Levene’s test, respectively. To evaluate the statistical significance of differences between two means, data were analyzed by Welch’s unpaired *t* test. To evaluate the statistical significance of differences among three or more means, data were analyzed by Tukey–Kramer test. When the variance differed among the groups, the Games–Howell test was used. Statistical tests were performed using JASP 0.19.1 software (JASP Team, 2024).

## Results

### *B. subtilis* strain YCN095 expressing the biofilm reporter P*tapA-mKate2*

The *B. subtilis* strain YCN095, which is based on the undomesticated strain NCIB3610, expresses the red fluorescent protein mKate2 under the control of the promoter for the biofilm matrix component gene *tapA* (P*tapA-mKate2*) (Norman et al., 2013; Gozzi et al., 2017). This strain formed robustly winkled colony biofilms with strong P*tapA-mKate2* expression on biofilm-inducing LBGM agar, compared to LB agar, over a period of 24 to 72 hours post-inoculation (**Figure 1A–B′**). Similarly, pellicle biofilms with robust wrinkles and strong P*tapA-mKate2* expression were observed on the surface of LBGM liquid during the same time frame, but not in LB liquid (**Figure 1C–D′**). These observations confirm that *B. subtilis* strain YCN095 functions well as a biofilm reporter strain.

**Figure 1.**
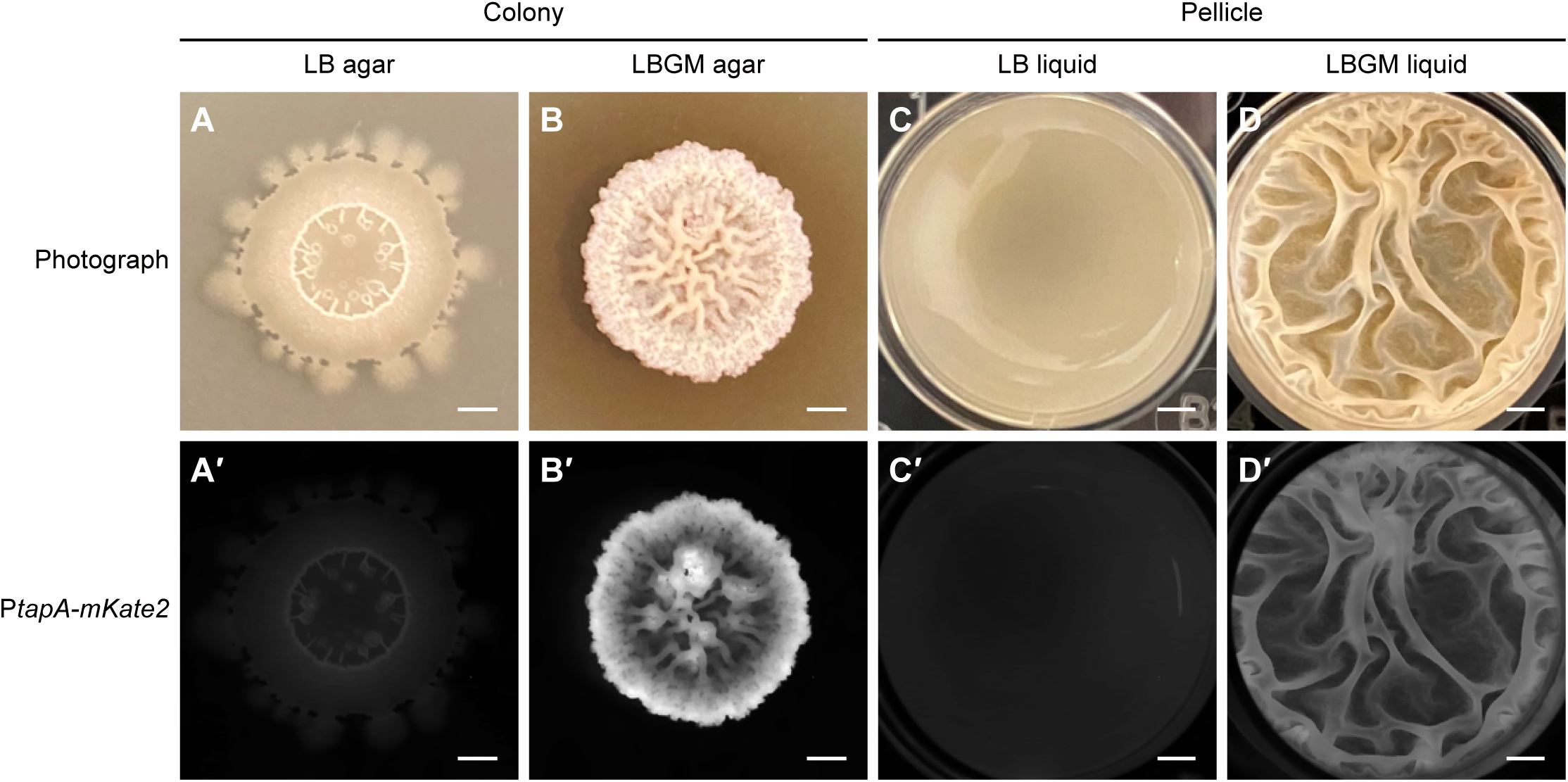
*B. subtilis* strain YCN095 forms robust wrinkled colony and pellicle biofilms with strong P*tapA-mKate2* expression. (A–D′) Representative photographs and red channel fluorescence images of colonies and pellicles formed 48 hours after inoculating LB and LBGM agar and liquid media with exponential phase YCN095 cultures are shown. YCN095 developed robust wrinkled colony biofilms (B, B′) and pellicle biofilms (D, D′) on LBGM agar and liquid, exhibiting strong P*tapA-mKate2* expression. Scale bars represent 2 mm.

### Biofilm formation and cell type composition in YCN095 lawns on NGM agar

To investigate biofilm formation and cell type composition on plate, YCN095 lawns were cultivated on NGM agar, a standard medium for rearing *C. elegans*. *B. subtilis* cell types were characterized under a 400x fluorescence upright microscope as follows (**Figure 2A**) (McKenney et al., 2013; Errington and Aart, 2020; Zhang et al., 2020). Motile cells were P*tapA-mKate2*-negative, rod-shaped, and approximately 2 µm in length. Matrix-producing cells were P*tapA-mKate2*-positive, non-motile, rod-shaped, and typically 2 µm or longer. Spores were small oval-shaped structures approximately 1 µm in length, with distinct bright and dark appearances at the surface and central focal planes, respectively. Sporulating cells were rod-shaped, approximately 2 µm in length, containing a single endospore.

**Figure 2.**
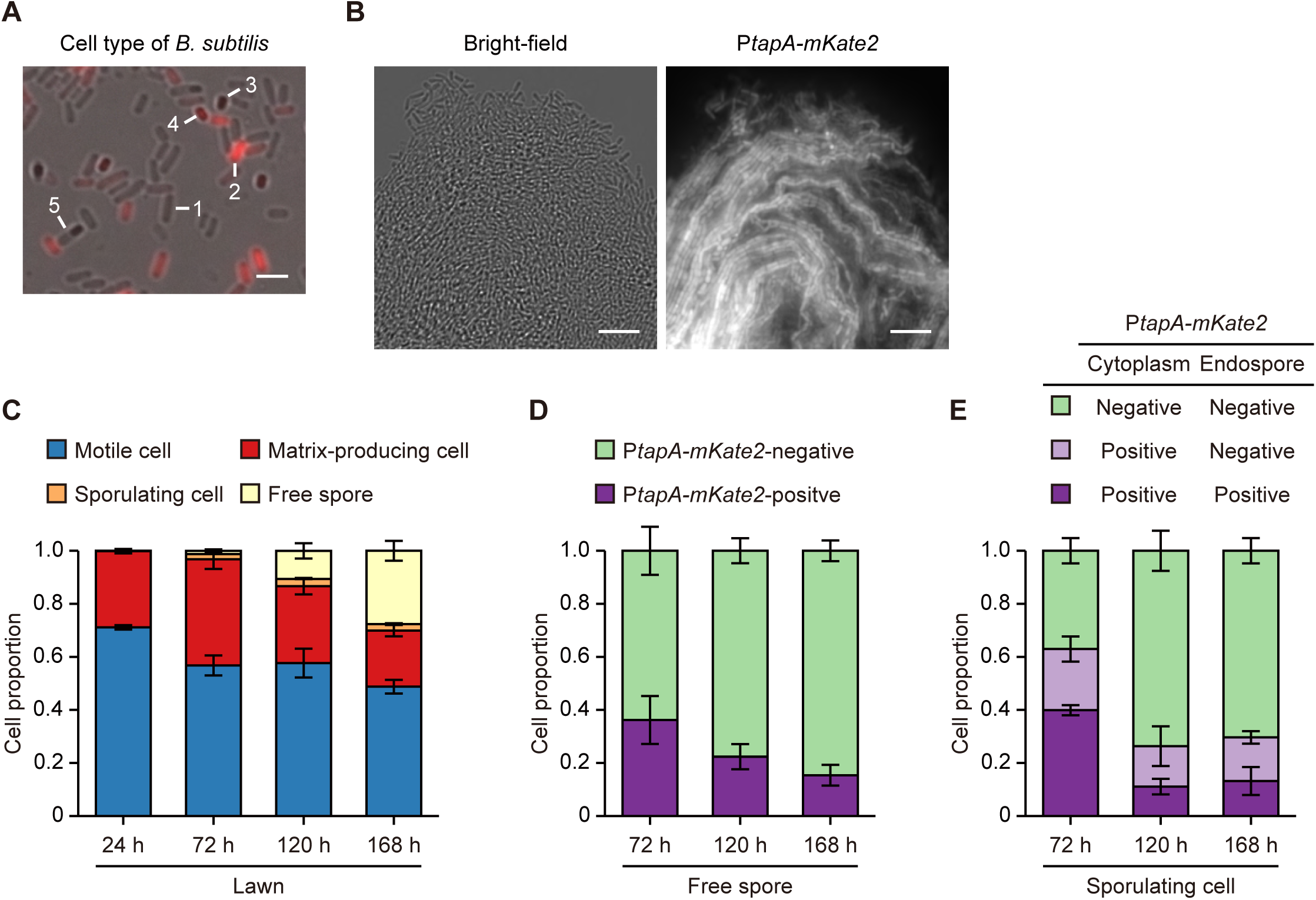
YCN095 lawns on NGM agar contain various cell types and biofilms. Exponential phase YCN095 cultures were inoculated onto NGM agar and incubated at 20°C for 168 hours to form lawns. (A) A representative merged bright-field and red channel image of a single-cell suspension from the lawn at 120 hours shows P*tapA-mKate2*-negative motile cells (1), P*tapA-mKate2*-positive matrix-producing cells (2), P*tapA-mKate2*-negative (3) and positive (4) free spores, and P*tapA-mKate2*-negative sporulating cells (5). Scale bar: 2 µm. (B) Bright-field and red channel images of a lawn sample scraped with a platinum wire at 120 hours show chain-forming and organized matrix-producing biofilms. The bright-field image also shows bright, refractive spores. Scale bars: 10 µm. (C) The composition of cell types in single-cell suspensions from lawns at 24, 72, 120, and 168 hours was determined. Data represent means ± SEM (*n* = 3 independent trials). Free spores were absent at 24 hours. Statistical comparisons were conducted among each cell type: motile cells (Turkey–Kramer test: 24 h vs. 168 h, *p* = 0.028); matrix-producing cells (Turkey–Kramer test: 72 h vs. 168 h, *p* = 0.015); sporulating cells (Games–Howell test: 24 h vs. 72 h and 120 h, *p* < 0.01; 24 h vs. 168 h, *p* = 0.015; 72 h vs. 168 h, *p* < 0.01); and free spores (Turkey–Kramer test: 72 h vs. 168 h, *p* < 0.01; 120 h vs. 168 h, *p* = 0.026). (D) Proportions of P*tapA-mKate2*-positive and negative free spores in lawns at 72, 120, and 168 hours are shown. Data represent means ± SEM (*n* = 3 independent trials). No significant differences were observed among P*tapA-mKate2*-positive free spores (Turkey–Kramer test). (E) Proportions of P*tapA-mKate2*-positive and negative sporulating cells in lawns at 72, 120, and 168 hours are shown. Data represent means ± SEM (n = 3 independent trials). The cell with P*tapA-mKate2*-negative cytoplasm but positive endospore was absent. Significant differences were detected among cells with P*tapA-mKate2*-positive cytoplasm and endospore (Turkey–Kramer test: 72 h vs. 120 h, *p* < 0.01; 72 h vs. 168 h, *p* = 0.013) and cells with P*tapA-mKate2*-negative cytoplasm and endospore (Turkey–Kramer test: 72 h vs. 120 h, *p* = 0.026; 72 h vs. 168 h, *p* = 0.038). No significant differences were found among cells with P*tapA-mKate2*-positive cytoplasm but negative endospore (Turkey–Kramer test).

YCN095 cells in the exponential phase were inoculated onto NGM agar and incubated at 20°C. The lawns did not display wrinkling appearance from 24 to 168 hours post-inoculation, unlike what was observed on LB or LBGM agar. Fluorescence microscopy revealed that matrix-producing cells were organized into chains and adhered tightly into three-dimensional structures, where spores were also observed, in the bacterial lawn samples scraped off from NGM plate with a platinum wire (**Figure 2B**). The results show that strain YCN095 forms biofilms on NGM agar.

The cell type composition of the YCN095 lawn was determined by analyzing single-cell suspensions of the whole lawn under a fluorescence upright microscope. As shown in **Figure 2C**, at 24 hours after inoculation, the lawn consisted mainly of motile cells and matrix-producing cells, with traces of sporulating cells and no free spores. Over time, the proportion of motile cells decreased, while the proportion of matrix-producing cells remained constant. The proportion of sporulating cells increased from 24 to 120 hours and plateaued thereafter. Free spores started to appear at 72 hours, increased over time, and reached 27.7% at 168 hours. These changes in cell type composition suggest that part of motility cells and matrix-producing cells differentiate into sporulating cells, which subsequently release spores during prolonged incubation.

Surprisingly, a fraction of free spores in the lawn were found to be P*tapA-mKate2*-positive (**Figure 2A, D**). Sporulating cells with P*tapA-mKate2* localized in both the cytoplasm and endospore were observed, with their proportion decreasing from 72 to 168 hours (**Figure 2E**). However, no sporulating cells were observed with P*tapA-mKate2* confined solely to the endospore (**Figure 2E**). These results suggest that P*tapA-mKate2* observed in the spores are potentially originated from the P*tapA-mKate2* present in the cytoplasm of sporulating cells, which may migrate into the endospore during sporulation, resulting in P*tapA-mKate2*-positive endospores.

### Biofilm formation not detected in the gut of *C. elegans* grown on YCN095 mixed-cell lawn

The ability of YCN095 to form biofilms in the gut of the *C. elegans* wild-type strain N2 was examined throughout the worm’s life cycle. Based on the cell type composition of the YCN095 lawn (**Figure 2C**), mixed-cell lawns at 5 to 8 days post-inoculation—containing motile cells, matrix-producing cells, sporulating cells, and free spores—were used as food. The guts of L4 larvae, young adults (day 2), middle-aged adults (day 7), and old adults (day 12) were analyzed using fluorescence microscopy.

At all developmental stages, single intact matrix-producing cells and P*tapA-mKate2*-positive spores were observed in the pharyngeal tract (**Figure 3A–A‴**), indicating that *C. elegans* ingests at least matrix-producing cells and spores from the lawn. However, aggregates of matrix-producing cells indicative of biofilm formation, even single matrix-producing cells, were absent in the intestinal tract (**Figure 3B–D‴**). Instead, diffused P*tapA-mKate2* fluorescence was observed throughout the intestinal tract (**Figure 3B″–D″**), suggesting that all the matrix-producing cells ingested by the worm were crushed by the pharyngeal grinder and resulted in the leaking of P*tapA-mKate2* into the intestinal tract. Furthermore, P*tapA-mKate2*-positive spores were observed drifting in the intestinal tract along with diffused P*tapA-mKate2* fluorescence (**Figure 3B″–D″, B‴–D‴; Supplementary Video**). Consistent with a previous report (Laaberki and Dworkin, 2008), the spores appeared to bypass the pharyngeal grinder due to their small size and reached the intestinal tract intact.

**Figure 3.**
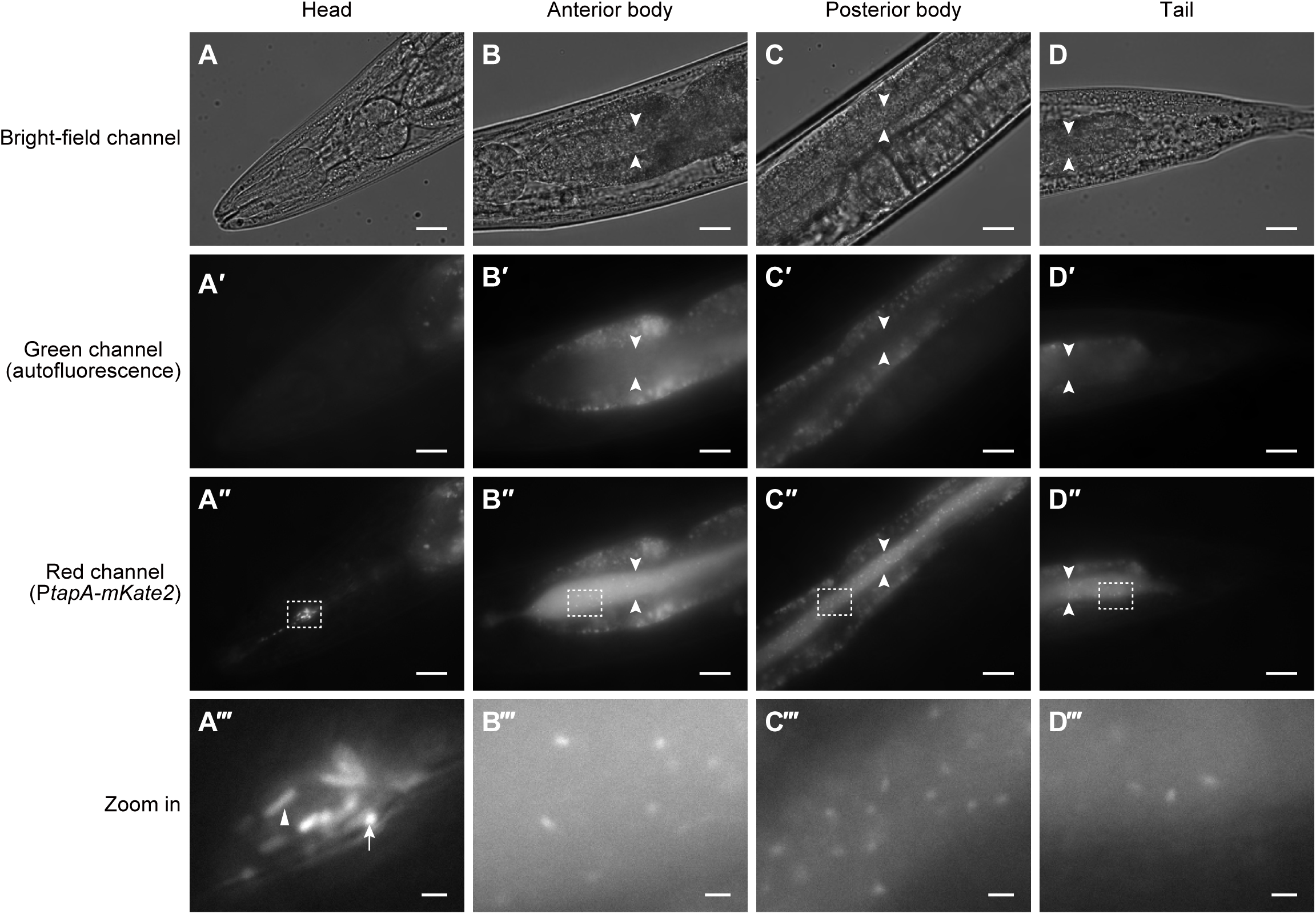
Biofilm is not formed in the gut of wild-type *C. elegans* with YCN095 mixed-cell lawns. (A–D‴) Representative bright-field and fluorescence channel images of the head, anterior body, posterior body, and tail regions of adult day 7 wild-type N2 worms grown on day 5–8 YCN095 mixed-cell lawns are shown. The white arrowheads in B–D″ indicate the borders of the intestinal lumen. Autofluorescence of intestinal cells is visible in the green channel (A′–D′) and co-localizes in the red channel (A″–D″). The specific fluorescence in the red channel originates from P*tapA-mKate2* (A″–D″). Enlarged views of regions outlined in A″–D″ are presented in A‴–D‴. (A–A‴) Rod-shaped matrix-producing cells positive for P*tapA-mKate2* (triangle) and smaller, oval-shaped P*tapA-mKate2*-positive spores (arrow) are visible in the pharyngeal tract. (B–D‴) In the intestinal tracts, neither aggregates of matrix-producing cells nor single cells were observed. Instead, diffuse P*tapA-mKate2* fluorescence and small, oval-shaped P*tapA-mKate2*-positive spores were observed. Scale bars represent 20 µm, except for enlarged views, which are 2 µm.

To corroborate these observations, food-switching experiments were conducted using *E. coli* OP50-GFP, which constitutively expresses GFP. Adult day 7 worms grown on the YCN095 mixed-cell lawn were transferred to the OP50-GFP lawn and cultured for 60 minutes. Fluorescence microscopy revealed that diffused P*tapA-mKate2* and P*tapA-mKate2*-positive spores were gradually expelled from the intestinal tract, starting within 10 minutes after the transfer (**Figure 4A–A″**). After 60 minutes, the intestinal lumen was filled exclusively with intact OP50-GFP cells and diffused GFP (**Figure 4B–B″**). There were no P*tapA-mKate2*-positive, matrix-producing cells attached to the intestinal apical membrane that should have been observed when biofilms formed in the intestinal tract (**Figure 4B′**). Conversely, adult day 7 worms grown on the OP50-GFP lawn were transferred to the YCN095 mixed-cell lawn and cultured for 60 minutes, a time period that is not sufficient for biofilm formation. OP50-GFP was progressively displaced by diffused P*tapA-mKate2* and P*tapA-mKate2*-positive spores (**Figure 4C–D″**), reproducing the intestinal tract images of worms grown on the YCN095 mixed-cell lawn alone. These findings indicate that *B. subtilis* YCN095 strain is unable to form biofilms in the guts of *C. elegans* wild-type N2 under a standard *C. elegans* culture condition on NGM plates.

**Figure 4.**
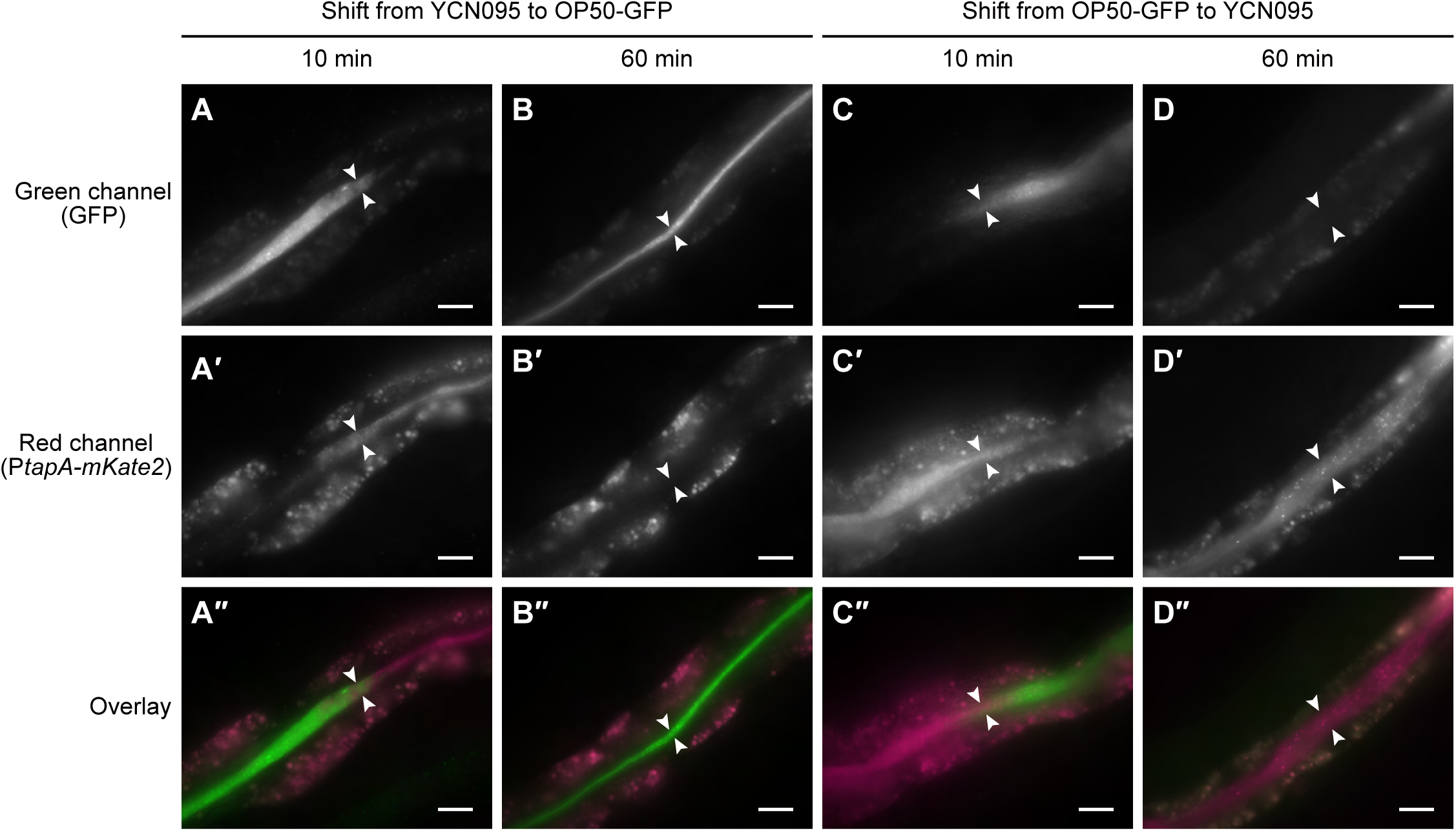
P*tapA-mKate2* fluorescence in the intestinal tracts of wild-type worms with YCN095 mixed-cell lawns is transient and derived from the diet. (A–D″) Wild-type N2 worms grown on day 5–8 YCN095 mixed-cell lawns or OP50-GFP lawns were transferred to OP50-GFP lawns or day-5 YCN095 mixed-cell lawns, respectively, and cultured for 10 or 60 minutes. Representative images of the posterior body of worms with heads oriented to the left are shown for fluorescence channels and merged views. The white arrowheads indicate the borders of the intestinal lumen. Fluorescence outside the intestinal tracts represents autofluorescence of intestinal cells. Upon food switching from YCN095 to OP50-GFP, diffuse P*tapA-mKate2* fluorescence, including P*tapA-mKate2*-positive spores in the intestinal tract, was pushed out by diffuse GFP containing intact OP50-GFP (A–A″) and completely disappeared after 60 minutes (B–B″). Conversely, upon switching from OP50-GFP to YCN095, diffuse GFP containing intact OP50-GFP in the intestinal tract was pushed out by diffuse P*tapA-mKate2* fluorescence, including P*tapA-mKate2*-positive spores (C–C″), and completely disappeared after 60 minutes (D–D″). Scale bars: 20 µm.

### Determination of total motile cells and spores in the gut of *C. elegans* grown on YCN095 mixed-cell lawn

Fluorescence microscopy can only confirm P*tapA-mKate2*-positive cells and spores in the *C. elegans* gut, while the bright-field images are too complex to deconvolute for determining the existence of P*tapA-mKate2*-negative cells and spores. Thus, we performed CFU assays to determine the presence of YCN095 total motile cells and spores in the gut of *C. elegans* wild-type N2 regardless of their fluorescence. Adult day 2 and day 7 worms grown on either the YCN095 mixed-cell lawn or the OP50 lawn were surface-sterilized, and CFU counts were performed on worm lysates containing only gut bacteria. For worms fed the YCN095 lawn, spore counts were determined based on the CFU obtained after heat treatment (80°C for 20 minutes), exploiting the heat resistance of spores. Since matrix-producing cells were absent in the intestinal tract (**Figure 3B–D‴**), the motile cell count was estimated by subtracting the CFU of spores from the total CFU of untreated lysates. For worms fed the OP50 lawn, the total bacterial load was estimated from untreated lysates.

OP50 cells were present in the gut of both day 2 and day 7 adults at levels consistent with previous studies (Portal-Celhay and Blaser, 2012; Gomez et al., 2012), with notable increases in bacterial load from day 2 to day 7, likely due to age-related declines in gut function (**Figure 5**). Similarly, both motile cells and spores were detected in worms fed the YCN095 lawn, with substantial increases in both cell types from day 2 to day 7 (**Figure 5**). The numbers of free spores in *C. elegans* gut were higher than or comparable to motile cells, while on the mixed plate motile cells always maintained the largest fraction (**Figure 2C**), likely due to the pharyngeal crushing. These observations confirm that spores ingested by the worms reach the intestinal tract intact and suggest that motile cells exist in the *C. elegans* intestinal tract but do not differentiate into matrix-producing cells, precluding *in vivo* biofilm formation in *C. elegans* gut.

**Figure 5.**
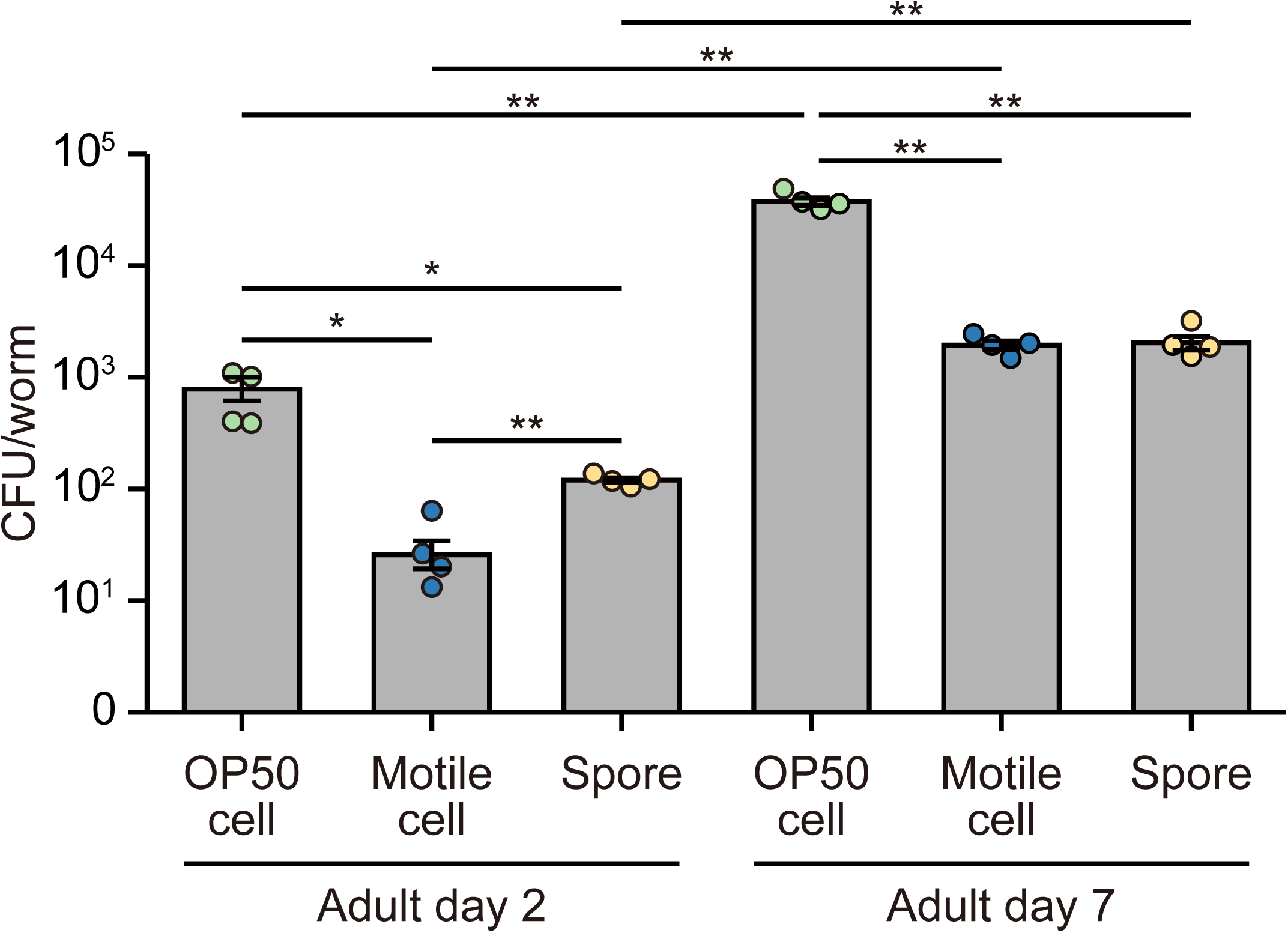
Motile cells and spores are present in the intestinal tracts of wild-type worms with YCN095 mixed-cell lawns. CFUs per worm were measured for wild-type N2 worms grown on OP50 lawns or day 5–8 YCN095 mixed-cell lawns until 2 or 7 days of adulthood. Data represent means ± SEM (*n* = 4 independent trials). \**p* < 0.05 and \*\**p* < 0.01 indicate significant differences between 2- and 7-day-old adults for the same bacterial cell type (Welch’s unpaired *t* test) or among bacterial cell types on adults of the same age (Games–Howell test).

### Biofilm formation not detected in the gut of *C. elegans* grown on YCN095 spore lawn

It has been reported that spores of the NCIB3610 strain can germinate and form biofilms in the intestinal tract of N2 worms when the worms are cultivated on a spore-only diet (Donato et al., 2017). This finding raises the possibility that differences in the cell type of *B. subtilis* ingested by *C. elegans* may contribute to the presence or absence of biofilm formation in the worm’s gut. To investigate this possibility, we fed purified spores of the YCN095 strain—prepared following the methodology described in Donato et al., 2017—to N2 worms and observed their intestinal tracts using fluorescence microscopy.

Purified spores were obtained by subjecting bacterial cultures rich in spores, cultivated in Schaeffer’s sporulation medium, to heat and lysozyme treatments. **Figure 6A** shows the morphology of free spores, germinated spores, and outgrown spores (He et al., 2018). Microscopic analysis confirmed that the purified spore suspension contained no motile cells, matrix-producing cells, or sporulating cells, with only a negligible presence of germinated and outgrown spores, achieving nearly 100% purity of free spores (**Figure 6B**). The purified spore suspension was subsequently applied onto NGM agar and allowed to dry, forming a dark brown lawn derived from the spores. This lawn was incubated at 20°C, and after 24 hours, it transformed into a grayish beige appearance, similar to that of mixed-cell lawns. Microscopic analysis of the entire lawn revealed that after 24 hours, the lawn contained free spores, germinated and outgrown spores, and motile cells in similar proportions (**Figure 6A, B**). By 48 hours, the population of motile cells had become predominant, with an increase in matrix-producing cells also observed (**Figure 6B**). The results suggest that free spores germinate on NGM agar, transitioning into motile cells, some of which subsequently differentiate into matrix-producing cells.

**Figure 6.**
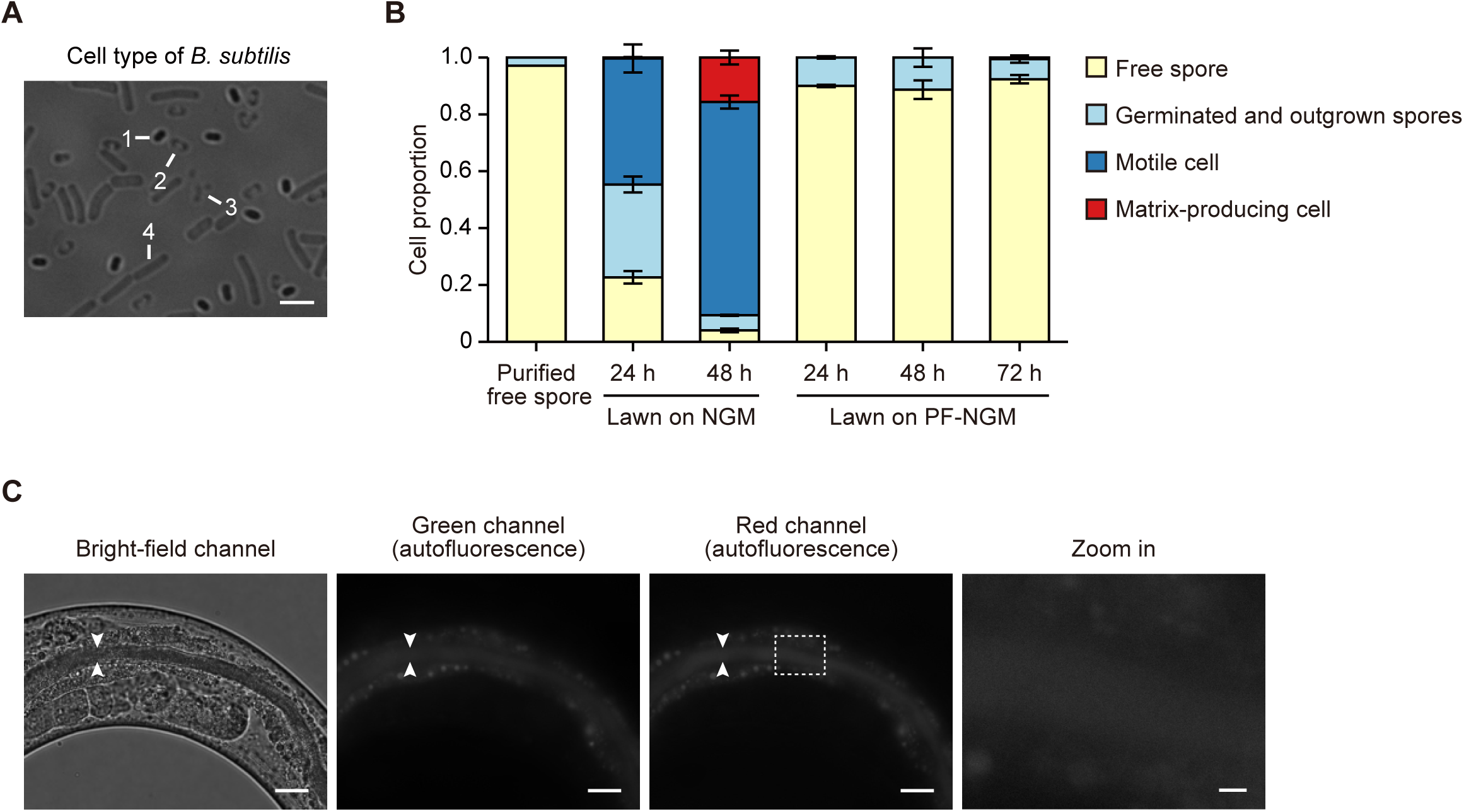
Biofilm is not formed in the gut of wild-type *C. elegans* with YCN095 spore lawns. (A) A representative bright-field image of free spores (1), germinated spores (2), outgrown spores (3), and motile cells (4) is shown. The image was derived from a single-cell suspension of a 24-hour NGM agar lawn. Scale bar: 2 µm. (B) Free spores were purified from liquid culture of YCN095 grown in Schaeffer’s sporulation medium, and their purity was confirmed. The purified spores were plated on NGM or PF-NGM agar, and the cell type composition of the lawns formed after 24, 48, and 72 hours was determined. Data represent mean ± SEM (*n* = 3 independent trials). On NGM agar, spores germinated and differentiated into motile cells and matrix-producing cells. On PF-NGM agar, most spores did not germinate, and the few spores that did germinate or outgrow remained in that state. Motile cells were observed only after 72 hours, and even then, at trace levels (0.6 ± 0.2%). No sporulating cells were observed on either NGM agar or PF-NGM agar. (C) Representative bright-field and fluorescence channel images of the posterior body of adult day 6 wild-type N2 worms grown on spore PF-NGM plates in a state within 48 hours of preparation are shown. The white arrowheads indicate the borders of the intestinal lumen. Autofluorescence of intestinal cells and intestinal tract is visible in the green channel and co-localizes in the red channel. Enlarged views of rectangular regions are presented. Neither aggregates of matrix-producing cells nor single cells were observed in the intestinal tract. Scale bars represent 20 µm, except for an enlarged view, which is 2 µm.

Consistent with earlier other studies (Laaberki and Dworkin, 2008; Goya et al., 2020), purified spores were also applied onto PF-NGM agar, dried, and cultured at 20°C for 3 days. The dark brown lawn remained unchanged in appearance throughout this period. Microscopic analysis revealed that the lawns after 24 and 48 hours consisted predominantly of free spores, with minor amounts of germinated and outgrown spores and no detectable motile, matrix-producing, or sporulating cells (**Figure 6B**). At 72 hours, while the proportions of free spores and germinated and outgrown spores remained largely unaltered, a trace amount (0.6%) of motile cells was observed (**Figure 6B**). Notably, all free spores, germinated spores, and outgrown spores, both immediately after purification and on PF-NGM agar, were P*tapA-mKate2* negative and exhibited weak autofluorescence, for reasons that remain unclear. The results suggest that nutrient-poor conditions on PF-NGM agar inhibit the germination of most free spores. Even in cases where limited germination occurs, the resultant cells fail to grow into motile cells.

The fact that spores did not geminate on PF-NGM provided an opportunity to evaluate whether spores can geminate in worm gut and form biofilm without interferences from germinated cells in the bacterial lawn on plate. As previously reported (Laaberki and Dworkin, 2008), N2 worms cultivated from L1 larvae on the spore lawn exhibited marked growth delays, likely attributable to the nutrient limitations imposed by the spores. To circumvent this issue, L1 larvae were initially grown on OP50-GFP lawns until reaching day 1 of adulthood, after which they were cultured on the spore lawn until day 6 of adulthood, with the lawns kept in a condition prepared within 48 hours. Microscopic analysis revealed that the intestinal tracts of adult day 6 worms were densely populated with spores, which were readily visible in the bright-field channel (**Figure 6C**). However, neither single matrix-producing cells nor aggregates indicative of biofilm formation were observed (**Figure 6C**). Only weak, dense autofluorescence was detectable, likely originating from the spores ingested by the worms (**Figure 6C**). Thus, we conclude that *B. subtilis* spores ingested by *C. elegans* do not lead to biofilm formation in the intestinal tract.

### Impact of life and death on gut biofilm formation in *C. elegans*

To determine whether differences in the internal environment between life and death influence biofilm formation, we examined biofilm formation in the gut of deceased *C. elegans*. Specifically, N2 worms at the old adult stage (adult day 16) reared on YCN095 mixed-cell lawns were cultured for an additional 2 days. Subsequently, worms that had succumbed to natural aging during this period were analyzed using a fluorescence upright microscope. In the intestinal tracts of living adult day 16 worms, P*tapA-mKate2*-positive spores were observed alongside autofluorescence associated with aging (**Figure 7A-A‴**). In contrast, in many worms that had died of old age, aggregates of matrix-producing cells were observed locally in the intestine, where the intestinal cells and intestinal tract were indistinguishable due to tissue decline caused by death, along with autofluorescence that had spread throughout the body (**Figure 7B–B‴**). Moreover, in many individuals, biofilm formation appeared to have advanced significantly, likely spreading from the intestine to the body cavity (**Figure 7C–C‴**). These findings suggest that specific functions associated with the life activities of *C. elegans* inhibit biofilm formation in the intestinal tract.

**Figure 7.**
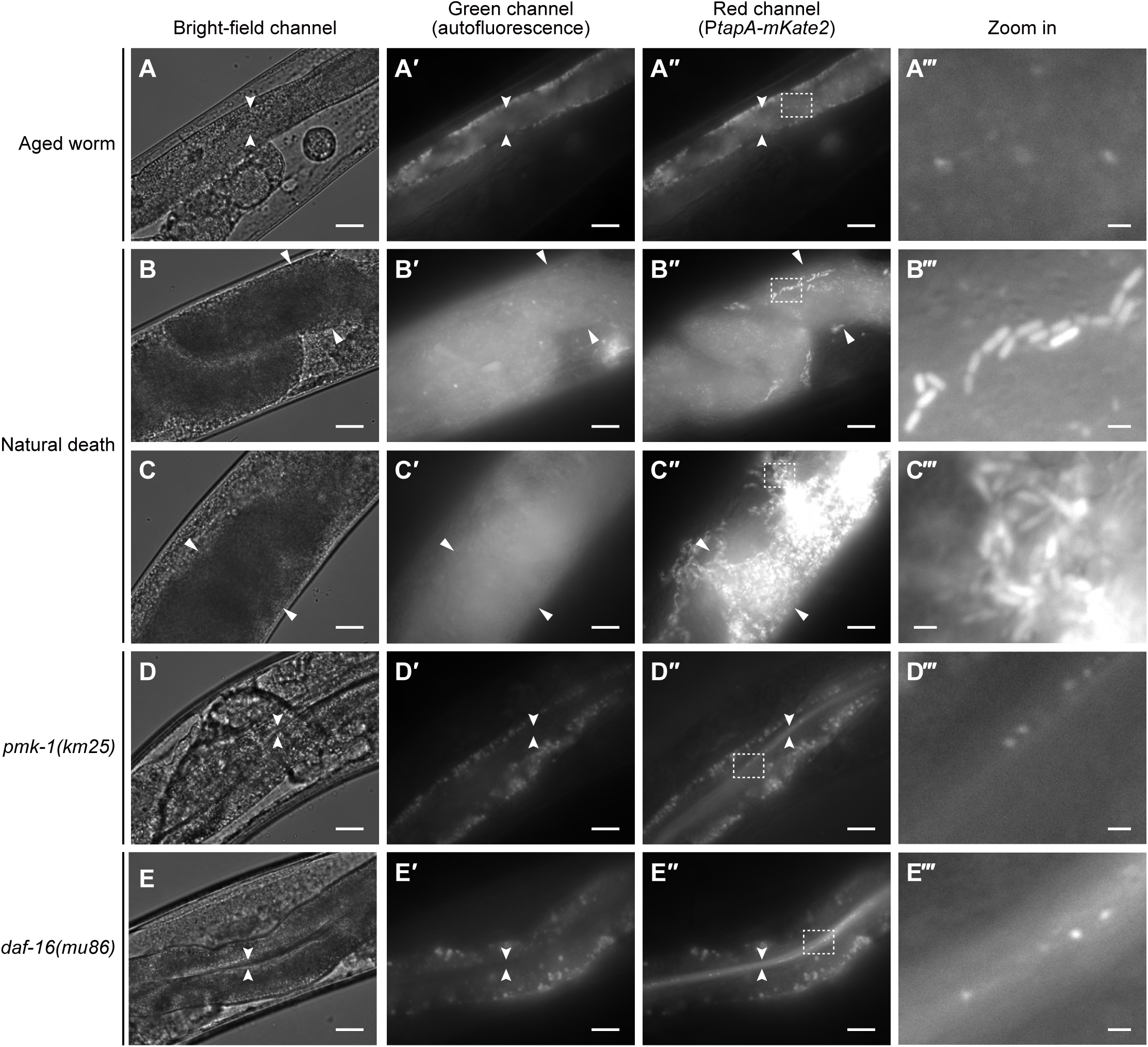
Biofilm formation in the intestinal tract is blocked by a life-dependent function of *C. elegans*. Representative bright-field and fluorescence images of the posterior body of live aged worms (A–A‴), worms that died of old age (B–C‴), and mutants with impaired innate immunity, *pmk-1(km25)* (D–D‴) and *daf-16(mu86)* (E–E‴), are shown. The white arrowheads in A–A″ and D–D″ indicate the boundaries of the intestinal lumen, while the triangles in B–C″ indicate the boundary of the intestine. Enlarged views of the areas outlined in A″–E″ are presented in A‴–E‴. (A–A‴) Wild-type N2 worms were cultured on YCN095 mixed-cell lawns until day 16 of adulthood. Autofluorescence from intestinal cells and the lumen was visible in the green channel (A′) and corresponded to the same regions in the red channel (A″). Neither aggregates of matrix-producing cells nor single cells were observed in the intestinal tract, but P*tapA-mKate2*-positive spores were observed (A″, A‴). (B–C‴) Wild-type N2 worms were cultured on YCN095 mixed-cell lawns until day 16 and observed after an additional 2 days when natural death occurred. Autofluorescence was detected throughout the body in the green channel (B′, C′), corresponding to regions in the red channel (B″, C″). The specific fluorescence in the red channel originated from P*tapA-mKate2*. Aggregates of matrix-producing cells, indicative of biofilm formation, were visible in swollen intestines where intestinal cell structures were no longer distinguishable (B″–C‴). Some individuals exhibited extensive biofilm formation spreading into the body cavity (C″, C‴). (D–E‴) *pmk-1(km25)* and *daf-16(mu86)* mutants were cultured on day 5-8 YCN095 mixed-cell lawns and observed on day 7 of adulthood. Autofluorescence of intestinal cells was visible in the green channel (D′, E′) and corresponded to regions in the red channel (D″, E″). The specific fluorescence in the red channel originated from P*tapA-mKate2*. No matrix-producing cells were observed in the intestinal tracts, but P*tapA-mKate2*-positive spores and diffuse P*tapA-mKate2* fluorescence were present (D″–E‴). Scale bars represent 20 µm, except for enlarged views, which are 2 µm.

The innate immune system in *C. elegans* is primarily regulated by the *pmk-1*/p38 MAPK kinase cascade and the *daf-16*/FOXO transcription factor activation pathway (McHugh et al., 2020). To determine whether these pathways influence biofilm formation in living worms, we analyzed *pmk-1(km25)* and *daf-16(mu86)* mutants at adult days 2, 7, and 12 grown on YCN095 mixed-cell lawns. In these mutants, no single cells or aggregates of matrix-producing cells were observed in the intestinal tracts (**Figure 7D–E‴**). Instead, P*tapA-mKate2*-positive spores and diffused P*tapA-mKate2* fluorescence were detected, resembling the observations in N2 worms (**Figure 7D–E‴**). These results indicate that the primary innate immune pathways in *C. elegans* are not directly involved in regulating biofilm formation in the intestinal tract. Instead, other life-dependent functions appear to play a critical role in regulating this process.

## Discussion

Biofilm-forming related pathways were shown to be critical for the beneficial effect from biofilm-forming *B. subtilis* NCIB3610 strain, including increased lifespan, enhanced resistance, and protection against neurotoxic agents (Donato et al., 2017; Smolentseva et al., 2017; Goya et al., 2020; Cogliati et al., 2020). However, it was not clear whether the beneficial effects were a result of live bacteria’s biofilm formation or the components from biofilm-forming bacteria. Previous reports showed contradictory results, where Donato et al. reported differential colonization and biofilm formation for biofilm-proficient and biofilm-deficient strains and Smolentseva et al. reported no differences in colonization (Donato et al., 2017; Smolentseva et al., 2017). In this study, we aimed to provide direct evidence in biofilm formation of *B. subtilis* NCIB3610 in *C. elegans* gut *in vivo*. To our surprise, our results showed no biofilm formation in *C. elegans* gut. Instead, free matrix-producing spores which were likely from food—both free spores that passed the pharyngeal grinder and sporulating cells that were grinded and released spores—were found freely “drifting” in the lumen instead of attached to gut surface, and the diffused fluorescence in the gut lumen indicated grinded cell contents instead of intact cells. Food switching experiments further confirmed that there were no matrix-producing cells/spores attached to the gut surface. In the below section, we will discuss our observations in detail, propose potential explanations for the discrepancy with previous reports, and suggest approaches for future studies.

The status of bacterial food fed to *C. elegans* is important for the results of worm-microbe interactions. Many studies have reported that *B. subtilis*, including the NCIB3610 strain (Sánchez-Blanco et al., 2016; Donato et al., 2017; Smolentseva et al., 2017; Goya et al., 2020; Cogliati et al., 2020) and other strains (Garsin et al., 2003; Gusarov et al., 2013; Iatsenko et al., 2014; Sánchez-Blanco et al., 2016; Donato et al., 2017; Hoang et al., 2019; Choi et al., 2020; Arvin et al., 2023), is a beneficial bacterium for *C. elegans*. However, even in experiments using the NCIB3610 strain which forms abundant biofilms, it was assumed that only motile cells or purified spores were consumed by *C. elegans*, leaving cell types such as the important biofilm matrix-producing cells unknown. Thus, we first clarified the cell type composition of the lawn of the YCN095 strain on *C. elegans* NGM agar. Our microscopic analysis revealed the presence of motile cells, matrix-producing cells, sporulating cells, and free spores, with notable changes in their proportions as the culture period progressed (**Figure 2C**). Specifically, it was suggested that motile cells and matrix-producing cells differentiated into sporulating cells, which subsequently formed spores (**Figure 2C**). Interestingly, although the wrinkles characteristic of *B. subtilis* biofilms were not observed in the YCN095 lawn on NGM plates, three-dimensional aggregates of matrix-producing cells, namely biofilms, were observed in samples scraped from lawns on NGM plates under microscope (**Figure 2B**). Most of the purified free spores on NGM agar germinated within 24 hours of inoculation to become motile cells (**Figure 6A, B**), followed by the appearance of matrix-producing cells after 48 hours (**Figure 6B**). In contrast, spore germination was suppressed on PF-NGM agar (**Figure 6B**), enabling the maintenance of a spore plate. A small amount of germinated and outgrown spores was present on the spore PF-NGM plate (**Figure 6B**) which can be a potential nutrient source. Our findings on the cell type composition in the lawn provide a solid foundation for subsequent *C. elegans* feeding experiments.

Next, we monitored biofilm formation in the gut of *C. elegans* throughout its lifespan using both day 5–8 YCN095 mixed-cell plates containing all *B. subtilis* cell types and spore PF-NGM plates. No aggregates of matrix-producing cells indicative of biofilm formation were observed in the gut of living worms from larvae to old adults (**Figures 3B–D‴, 6C, 7A–A‴**). Food switching experiments showed that the diffused fluorescence and spores with fluorescence can be readily replaced by *E. coli* food with no indication of gut surface attachment, another indication of biofilm formation (**Figure 4A–B″**). CFU assays confirmed the presence of motile cells in the gut (**Figure 5**), suggesting that differentiation into matrix-producing cells was inhibited, thereby preventing biofilm formation. In contrast, biofilm formation was observed in the intestines of worms that had died of old age (**Figure 7B–C‴**), indicating that life-dependent factors strongly influence the lack of biofilm formation in the intestinal tract. Two pathways related to innate immunity, *pmk-1* and *daf-16*, in *C. elegans* were tested by *C. elegans* mutants and showed identical results like the N2 worms (**Figure 7D–E‴**). Other possible factor inhibiting biofilm formation in the intestinal tract is its acidic pH. While *B. subtilis* forms biofilms at pH 5.0–9.5, with an optimal pH of 7.4–8.0 (Sharipova et al., 2023), the mean pH of the *C. elegans* intestinal lumen is 3.9 (Chauhan et al., 2013), which may suppress biofilm formation. In fact, after death, the intestinal pH increases to approximately 6.0, as it reflects the surrounding medium (Chauhan et al., 2013). Another potential factor is the strict regulation of biofilm-inducing elements such as iron and manganese by the SMF protein, a divalent metal transporter ortholog (Au et al., 2009). This regulation likely maintains metal ion concentrations at low levels, unfavorable for biofilm formation. Upon death, homeostasis is disrupted, leading to increased concentrations of metal ions from the surrounding medium, which may promote biofilm formation. Thus, the gut environment of living *C. elegans* appears unsuitable for biofilm formation. In nature, *B. subtilis* interacts with various soil bacteria and fungi, which may influence processes like biofilm formation, sporulation, and motility (Andrić et al., 2020). However, under the culture conditions of this study, such interactions were absent, potentially suppressing *B. subtilis* biofilm formation in the *C. elegans* gut.

Our results indicate that all matrix-producing cells of the YCN095 strain ingested by *C. elegans* are crushed by the pharyngeal grinder, releasing their contents into the intestinal lumen (**Figure 3**). Furthermore, the ingested spores, likely due to their small sizes, appear to pass through the pharyngeal grinder and reach the intestinal tract (**Figure 3**). Similarly, intact motile cells, which are approximately of similar size as matrix-producing cells, are also presumed to be crushed by the pharyngeal grinder and do not reach the intestinal lumen. Based on this assumption, the presence of motile cells confirmed by the CFU assay (**Figure 5**) is likely a result of the germination of free spores in the intestinal tract. Spores geminated into motile cells in the intestinal lumen but not into matrix-producing cells, indicated by the lack of P*tapA* reporter fluorescence. The increase in motile cells observed in middle-aged worms (**Figure 5**) may be attributed to the accumulation of free spores, followed by their germination into motile cells, as a consequence of age-related decline in gut function. It should be noted that in this CFU assay, we did not perform the step of incubating collected worms in a buffer for a certain period to remove transient bacteria in the intestinal lumen, in order to prevent the expulsion of free spores from the gut. Therefore, it is currently unclear whether the motile cells detected in this assay are colonized in the intestine or not. Future study may employ *B. subtilis* with a reporter specific to motile cells to visualize potential colonization of motile cells to the intestinal apical membrane.

As previously mentioned, two research groups reported that *B. subtilis* strain NCIB3610 forms or exists as a biofilm in the intestinal tract of *C. elegans* (Donato et al., 2017; Smolentseva et al., 2017), which contradicts our results. Such discrepancy may be explained by the nature of relevant protocols used. Previous studies assessed biofilm formation based on fluorescence intensity in intestinal lumen from biofilm-related reporters without differentiating amongst diffused fluorescence, free spores or matrix-producing cell aggregates. Based on our results, the observed fluorescence in the intestinal tract in previous studies may have originated from free spores and fluorescent proteins leaked from matrix-producing cells of dietary origin that were crushed by the pharyngeal grinder, as Smolentseva et al. used mixed-cell plates for feeding and Donato et al. used purified spore on NGM plates which were plausibly de facto mixed-cell plates during incubation (**Figure 6B**). Thus, echoing and expanding what was suggested by Smolentseva et al., colonization and subsequent biofilm formation may not be the mechanisms for the beneficial effects of biofilm-forming *B. subtilis* to *C. elegans*, but rather the components from matrix-producing biofilm cells that can elicit certain host responses. Finally, we would also like to note that significant genetic variations exist among *B. subtilis* strains used in different laboratories worldwide (Zeigler et al., 2008; Patrick and Kearns, 2009; Mcloon et al., 2011). Thus, the possibility that differences in biofilm formation in the intestinal tract were caused by mutations in the NCIB3610 strain cannot be completely ruled out.

As described above, feeding on the biofilm-forming *B. subtilis* strain NCIB3610 has been reported to confer extended lifespan, resistance to various stresses and pathogenic infections, and protection against α-synuclein and amyloid-β-associated toxicity (Donato et al., 2017; Smolentseva et al., 2017; Goya et al., 2020; Cogliati et al., 2020). These effects are abrogated when worms are fed on isogenic biofilm-deficient mutant strains. Nitric oxide (NO) and quorum-sensing pentapeptides (CSF) derived from the *B. subtilis* biofilm have been suggested as the factors responsible for these beneficial effects (Donato et al., 2017; Goya et al., 2020; Cogliati et al., 2020). Our results, as discussed above, indicate that all matrix-producing cells consumed from mixed-cell lawns are crushed by the pharyngeal grinder and that biofilm does not form in the intestinal tract (**Figure 3**). Matrix-producing cells in mixed-cell lawns likely contain abundant NO and CSF because *B. subtilis* cultured under biofilm-forming conditions *in vitro* produces these molecules in high quantities (Donato et al., 2017). It is plausible that NO and CSF released from matrix-producing cells lysed by the pharyngeal grinder diffuse and act in the intestinal tract, leading to these beneficial effects. Supporting this possibility, the lifespan-extending effects can be achieved by feeding with lysates instead of bacterial lawns (Saito et al., 2023), and the protective effects against α-synuclein-associated toxicity can be achieved through exogenous addition of NO and CSF (Goya et al., 2020). Testing our hypothesis that the contents derived from matrix-producing cells in the diet are responsible for these beneficial effects can be accomplished by comparing lysates of biofilm-forming and biofilm-deficient mutant strains as dietary sources. Identifying the differing components within these lysates may provide useful insights into the mechanisms underlying these effects.

## Ethics

This work did not require ethical approval from a human subject or animal welfare committee.

## Declaration of AI use

We have not used AI-assisted technologies in creating this article.

## Authors’ contributions

Y.S.: conceptualization, methodology, investigation, formal analysis, validation, data curation, visualization, project administration, writing-original draft, writing-review & editing. T.A.Q.: conceptualization, methodology, funding acquisition, project administration, resources, supervision, writing-review & editing.

## Acknowledgments

We thank Prof. Yunrong Chai (Northeastern University, Boston, USA) for generously gifting *B. subtilis* strains. Some *C. elegans* and *E. coli* strains were kindly supplied by the Caenorhabditis Genetics Center (University of Minnesota, Minneapolis, USA), which is funded by the National Institutes of Health, National Center for Research Resources. Open access fee was supported by Michigan State University. We thank Prof. Lee R. Kroos at Michigan State University for his helpful consultation during project conceptualization. We thank Prof. Kin Sing Lee and Jennifer Hinman at Michigan State University for their useful discussion on biofilm formation associated with aging. This study was conducted during the author’s study abroad as a research fellow (Yasuaki Saitoh), supported by the employment at Kitasato University, which provided continuous salary support.

## References

Andrić S, Meyer T, Ongena M. 2020 *Bacillus* responses to plant-associated fungal and bacterial communities. Front. Microbiol. 11, 1350. (doi:10.3389/fmicb.2020.01350).

Arata Y, Oshima T, Ikeda Y, Kimura H, Sako Y. 2020 OP50, a bacterial strain conventionally used as food for laboratory maintenance of *C. elegans*, is a biofilm formation defective mutant. MicroPubl. Biol. 10.17912/micropub.biology.000216. (doi:10.17912/micropub.biology.000216).

Arnaouteli S, Bamford NC, Stanley-Wall NR, Kovács ÁT. 2021 *Bacillus subtilis* biofilm formation and social interactions. Nat. Rev. Microbiol. 19, 600–614. (doi:10.1038/s41579-021-00540-9).

Arvin CL, Sibila Z, Lamendella R, Chan J, Staab T. 2023 The impact of *C. elegans* ceramide glucosyltransferase enzymes on the beneficial effects of *B. subtilis* lifespan. MicroPubl. Biol. 10.17912/micropub.biology.000758. (doi:10.17912/micropub.biology.000758).

Au C, Benedetto A, Anderson J, Labrousse A, Erikson K, Ewbank JJ, Aschner M. 2009 SMF-1, SMF-2 and SMF-3 DMT1 orthologues regulate and are regulated differentially by manganese levels in *C. elegans*. PLoS One 4, e7792. (doi:10.1371/journal.pone.0007792).

Brenner S. 1974 The genetics of *Caenorhabditis elegans*. Genetics 77, 71–94. (doi:10.1093/genetics/77.1.71).

Buret AG, Allain T. 2023 Gut microbiota biofilms: From regulatory mechanisms to therapeutic targets. J. Exp. Med. 220, e20221743. (doi:10.1084/jem.20221743).

C. elegans Sequencing Consortium. 1998 Genome sequence of the nematode *C. elegans*: a platform for investigating biology. Science 282, 2012–2018. (doi:10.1126/science.282.5396.2012).

Chauhan VM, Orsi G, Brown A, Pritchard DI, Aylott JW. 2013 Mapping the pharyngeal and intestinal pH of *Caenorhabditis elegans* and real-time luminal pH oscillations using extended dynamic range pH-sensitive nanosensors. ACS Nano 7, 5577–5587. (doi:10.1021/nn401856u).

Choi HJ, Shin D, Shin M, Yun B, Kang M, Yang HJ, Jeong DY, Kim Y, Oh S. 2020 Comparative genomic and functional evaluations of *Bacillus subtilis* newly isolated from Korean traditional fermented foods. Foods 9, 1805. (doi:10.3390/foods9121805).

Cogliati S, Clementi V, Francisco M, Crespo C, Argañaraz F, Grau R. 2020 *Bacillus Subtilis* delays neurodegeneration and behavioral impairment in the Alzheimer’s disease model *Caenorhabditis elegans*. J. Alzheimers Dis. 73, 1035–1052. (doi:10.3233/JAD-190837).

Dirksen P, Assié A, Zimmermann J, Zhang F, Tietje AM, Marsh SA, Félix MA, Shapira M, Kaleta C, Schulenburg H, Samuel BS. 2020 CeMbio - The *Caenorhabditis elegans* Microbiome Resource. G3 (Bethesda) 10, 3025–3039. (doi:10.1534/g3.120.401309).

Donato V, Ayala FR, Cogliati S, Bauman C, Costa JG, Leñini C, Grau R. 2017 *Bacillus subtilis* biofilm extends *Caenorhabditis elegans* longevity through downregulation of the insulin-like signalling pathway. Nat. Commun. 8, 14332. (doi:10.1038/ncomms14332).

Errington J, Aart LTV. 2020 Microbe Profile: *Bacillus subtilis*: model organism for cellular development, and industrial workhorse. Microbiology 166, 425–427. (doi:10.1099/mic.0.000922).

Félix MA, Duveau F. 2012 Population dynamics and habitat sharing of natural populations of *Caenorhabditis elegans* and *C. briggsae*. BMC Biol. 10, 59. (doi:10.1186/1741-7007-10-59)..

Garsin DA, Villanueva JM, Begun J, Kim DH, Sifri CD, Calderwood SB, Ruvkun G, Ausubel FM. 2003 Long-lived *C. elegans daf-2* mutants are resistant to bacterial pathogens. Science 300, 1921. (doi:10.1126/science.1080147).

Gomez F, Monsalve GC, Tse V, Saiki R, Weng E, Lee L, Srinivasan C, Frand AR, Clarke CF. 2012 Delayed accumulation of intestinal coliform bacteria enhances life span and stress resistance in *Caenorhabditis elegans* fed respiratory deficient *E. coli*. BMC Microbiol. 12, 300. (doi:10.1186/1471-2180-12-300).

Goya ME, Xue F, Sampedro-Torres-Quevedo C, Arnaouteli S, Riquelme-Dominguez L, Romanowski A, Brydon J, Ball KL, Stanley-Wall NR, Doitsidou M. 2020 Probiotic *Bacillus subtilis* protects against α-synuclein aggregation in *C. elegans*. Cell Rep. 30, 367–380.e7. (doi:10.1016/j.celrep.2019.12.078).

Gozzi K, Ching C, Paruthiyil S, Zhao Y, Godoy-Carter V, Chai Y. 2017 *Bacillus subtilis* utilizes the DNA damage response to manage multicellular development. NPJ Biofilms Microbiomes 3, 8. (doi:10.1038/s41522-017-0016-3).

Gusarov I, Gautier L, Smolentseva O, Shamovsky I, Eremina S, Mironov A, Nudler E. 2013 Bacterial nitric oxide extends the lifespan of *C. elegans*. Cell 152, 818–830. (doi:10.1016/j.cell.2012.12.043).

Hashem A, Tabassum B, Fathi Abd Allah E. 2019 *Bacillus subtilis*: A plant-growth promoting rhizobacterium that also impacts biotic stress. Saudi J. Biol. Sci. 26, 1291–1297. (doi:10.1016/j.sjbs.2019.05.004).

He L, Chen Z, Wang S, Wu M, Setlow P, Li YQ. 2018 Germination, outgrowth, and vegetative-growth kinetics of dry-heat-treated individual spores of *Bacillus* species. Appl. Environ. Microbiol. 84, e02618–17. (doi:10.1128/AEM.02618-17).

Hoang KL, Gerardo NM, Morran LT. 2019 The effects of *Bacillus subtilis* on *Caenorhabditis elegans* fitness after heat stress. Ecol. Evol. 9, 3491–3499. (doi:10.1002/ece3.4983).

Iatsenko I, Yim JJ, Schroeder FC, Sommer RJ. 2014 *B. subtilis* GS67 protects *C. elegans* from Gram-positive pathogens via fengycin-mediated microbial antagonism. Curr. Biol. 24, 2720–2727. (doi:10.1016/j.cub.2014.09.055).

JASP Team. 2024 JASP (Version 0.19.1) [Computer software].

Kiontke KC, Félix MA, Ailion M, Rockman MV, Braendle C, Pénigault JB, Fitch DH. 2011 A phylogeny and molecular barcodes for *Caenorhabditis*, with numerous new species from rotting fruits. BMC Evol. Biol. 11, 339. (doi:10.1186/1471-2148-11-339).

Laaberki MH, Dworkin J. 2008 Role of spore coat proteins in the resistance of *Bacillus subtilis* spores to *Caenorhabditis elegans* predation. J. Bacteriol. 190, 6197–6203. (doi:10.1128/JB.00623-08).

McGhee JD. 2007 The *C. elegans* intestine. WormBook 6197–6203. (doi:10.1895/wormbook.1.133.1).

McHugh DR, Koumis E, Jacob P, Goldfarb J, Schlaubitz-Garcia M, Bennani S, Regan P, Patel P, Youngman MJ. 2020 DAF-16 and SMK-1 contribute to innate immunity during adulthood in *Caenorhabditis elegans*. G3 (Bethesda) 10, 1521–1539. (doi:10.1534/g3.120.401166).

McKenney PT, Driks A, Eichenberger P. 2013 The *Bacillus subtilis* endospore: assembly and functions of the multilayered coat. Nat. Rev. Microbiol. 11, 33–44. (doi:10.1038/nrmicro2921).

McLoon AL, Guttenplan SB, Kearns DB, Kolter R, Losick R. 2011 Tracing the domestication of a biofilm-forming bacterium. J. Bacteriol. 193, 2027–2034. (doi:10.1128/JB.01542-10).

Mielich-Süss B, Lopez D. 2015 Molecular mechanisms involved in *Bacillus subtilis* biofilm formation. Environ. Microbiol. 17, 555–565. (doi:10.1111/1462-2920.12527).

Norman TM, Lord ND, Paulsson J, Losick R. 2013 Memory and modularity in cell-fate decision making. Nature 503, 481–486. (doi:10.1038/nature12804).

Patrick JE, Kearns DB. 2009 Laboratory strains of *Bacillus subtilis* do not exhibit swarming motility. J. Bacteriol. 191, 7129–7133. (doi:10.1128/JB.00905-09).

Portal-Celhay C, Blaser MJ. 2012 Competition and resilience between founder and introduced bacteria in the *Caenorhabditis elegans* gut. Infect. Immun. 80, 1288–1299. (doi:10.1128/IAI.05522-11).

Saito R, Sato N, Okino Y, Wang DS, Seo G. 2023 *Bacillus subtilis* TO-A extends the lifespan of *Caenorhabditis elegans*. Biosci. Microbiota Food Health 42, 124–130. (doi:10.12938/bmfh.2022-057).

Sánchez-Blanco A, Rodríguez-Matellán A, González-Paramás A, González-Manzano S, Kim SK, Mollinedo F. 2016 Dietary and microbiome factors determine longevity in *Caenorhabditis elegans*. Aging (Albany NY) 8, 1513–1539. (doi:10.18632/aging.101008).

Schoenborn AA, Yannarell SM, Wallace ED, Clapper H, Weinstein IC, Shank EA. 2021 Defining the expression, production, and signaling roles of specialized metabolites during *Bacillus subtilis* differentiation. J. Bacteriol. 203, e0033721. (doi:10.1128/JB.00337-21).

Sharipova M, Rudakova N, Mardanova A, Evtugyn V, Akosah Y, Danilova I, Suleimanova A. 2023 Biofilm formation by mutant strains of bacilli under different stress conditions. Microorganisms 11, 1486. (doi:10.3390/microorganisms11061486).

Shineh G, Mobaraki M, Perves Bappy MJ, Mills DK. 2023 Biofilm formation, and related impacts on healthcare, food processing and packaging, industrial manufacturing, marine industries, and sanitation–a review. Appl. Microbiol. 3, 629–665. (10.3390/applmicrobiol3030044)

Siala A, Hill IR, Gray TRG. 1974 Populations of spore-forming bacteria in an acid forest soil, with special reference to *Bacillus subtilis*. Microbiol. 81, 183–190. (doi:10.1099/00221287-81-1-183).

Smolentseva O, Gusarov I, Gautier L, Shamovsky I, DeFrancesco AS, Losick R, Nudler E. 2017 Mechanism of biofilm-mediated stress resistance and lifespan extension in *C. elegans*. Sci. Rep. 7, 7137. (doi:10.1038/s41598-017-07222-8).

Vega NM, Gore J. 2017 Stochastic assembly produces heterogeneous communities in the *Caenorhabditis elegans* intestine. PLoS Biol. 15, e2000633. (doi:10.1371/journal.pbio.2000633).

Vlamakis H, Chai Y, Beauregard P, Losick R, Kolter R. 2013 Sticking together: building a biofilm the *Bacillus subtilis* way. Nat. Rev. Microbiol. 11, 157–168. (doi:10.1038/nrmicro2960).

Zeigler DR, Prágai Z, Rodriguez S, Chevreux B, Muffler A, Albert T, Bai R, Wyss M, Perkins JB. 2008 The origins of 168, W23, and other *Bacillus subtilis* legacy strains. J. Bacteriol. 190, 6983–6995. (doi:10.1128/JB.00722-08).

Zhang X, AI-Dossary A, Hussain M, Setlow P, Li J. 2020 Applications of *Bacillus subtilis* spores in biotechnology and advanced materials. Appl. Environ. Microbiol. 86, e01096–20. (doi:10.1128/AEM.01096-20).

